# Nicotinamide Riboside supplementation does not alter whole-body or skeletal muscle metabolic responses to a single bout of endurance exercise

**DOI:** 10.1101/2020.06.23.143446

**Authors:** Ben Stocks, Stephen P. Ashcroft, Sophie Joanisse, Yasir S. Elhassan, Gareth G. Lavery, Linda C. Dansereau, Ashleigh M. Philp, Gareth A. Wallis, Andrew Philp

## Abstract

Oral supplementation of the NAD^+^ precursor Nicotinamide Riboside (NR) has been reported to increase Sirtuin (SIRT) signalling, mitochondrial biogenesis and endurance capacity in rodent skeletal muscle. However, whether NR supplementation can elicit a similar response in human skeletal muscle is unclear. This study aimed to assess the effect of 7-day NR supplementation on exercise-induced transduction and transcriptional responses in skeletal muscle of young, healthy, recreationally active human volunteers. In a double-blinded, randomised, counter-balanced, crossover design, eight male participants (age: 23 ± 4 years, VO_2_peak: 46.5 ± 4.4 mL·kg^-1^·min^-1^) received one week of NR or cellulose placebo (PLA) supplementation (1000 mg·d^-1^) before performing one hour of cycling at 60% Wmax. Muscle biopsies were collected prior to supplementation and pre-, immediately and three-hours post-exercise from the medial vastus lateralis, whilst venous blood samples were collected throughout the trial. Global acetylation, auto-PARylation of PARP1, acetylation of p53^Lys382^ and MnSOD^Lys122^ were unaffected by NR supplementation or exercise. Exercise led to an increase in AMPK^Thr172^ (1.6-fold), and ACC^Ser79^ (4-fold) phosphorylation, in addition to an increase in PGC-1α (∼5-fold) and PDK4 (∼10-fold) mRNA expression, however NR had no additional effect on this response. There was also no effect of NR supplementation on substrate utilisation at rest or during exercise or on skeletal muscle mitochondrial respiration. Finally, NR supplementation blunted the exercise induced activation of skeletal muscle NNMT mRNA expression, but had no effect on mRNA expression of NMRK1, NAMPT or NMNAT1, which were not significantly affected by NR supplementation or exercise. In summary, one week of NR supplementation does not augment skeletal muscle signal transduction pathways implicated in mitochondrial adaptation to endurance exercise.

## Introduction

Nicotinamide adenine dinucleotide (NAD^+^), including its reduced form NADH, is a redox co-enzyme that shuttles hydride ions between processes of fuel oxidation, as well as within biosynthethic pathways [1]. In addition to central roles in these critical metabolic processes, NAD^+^ has emerged as a signalling moiety and an obligatory co-substrate for sirtuins (SIRTs), poly ADP-ribose polymerases (PARPs) and cyclic ADP-ribose synthetases [2]. Thus NAD^+^ is an important substrate in pathways governing metabolic adaptations, DNA repair and apoptosis [1, 2]. Given the regulatory role of NAD^+^ in lifespan extending processes, it is unsurprising that strategies to elevate cellular NAD^+^ are considered as promising therapies. For example, elevating cellular NAD^+^ *in vivo* leads to positive outcomes in murine models of diabetes [3, 4], ageing [3, 5, 6], obesity [7], vascular dysfunction [5], muscular dystrophy [8] and Alzheimer’s disease [9].

The vitamin B3 molecule nicotinamide riboside (NR) has emerged as one dietary strategy to elevate NAD^+^ *in vivo*. In rodents, oral NR supplementation increases fat oxidation (at least during the light, inactive phase) [7], promotes metabolic flexibility [10], improves insulin sensitivity and may improve endurance performance [7], although a trend towards impaired endurance performance has also been noted [11]. Mechanistically, NR supplementation increases SIRT1 and SIRT3 activity, deacetylation of peroxisome proliferator-activated receptor-γ coactivator 1-α (PGC-1α) and induces mitochondrial biogenesis [7, 8, 12, 13]. Interestingly, and somewhat unsurprisingly, the effects of NR supplementation are much more pronounced during models of elevated cellular stress [7, 8, 12-14].

Studies investigating NR supplementation in humans are in their infancy [6, 15-22]. Importantly, NR displays excellent safety and oral bioavailability in humans [6, 16, 17, 22], with NR supplementation reported to improve blood pressure [6], liver health [16] and physical function in the elderly [16], although the latter is not a consistent finding [6]. However, despite promising evidence from pre-clinical models [7, 10], several studies have reported no effect of chronic NR supplementation on mitochondrial volume, mitochondrial respiration, insulin sensitivity, body composition, cardiac function, lipolysis, VO_2_peak and resting or exercising substrate utilisation [6, 17-21], although one study has found increases in relative fat free mass and sleeping metabolic rate [21]. The effect of NR on mitochondrial biogenic signalling in skeletal muscle following exercise remains unstudied.

Therefore, the purpose of this study was to investigate the effects of oral NR supplementation on whole body substrate utilisation and skeletal muscle mitochondrial biogenic signalling at rest and following acute steady-state exercise in humans. It was hypothesised that NR supplementation would increase whole body fat oxidation during exercise and augment SIRT1, SIRT3 and PGC-1α signalling in the post-exercise period compared to placebo.

## Materials and Methods

### Participants

Eight recreationally active males (mean ± SD: age, 23 ± 4 years; body mass, 72.4 ± 5.3 kg; peak oxygen uptake (VO_2_peak), 46.5 ± 4.4 mL·kg^-1^·min^-1^; maximal aerobic power (Wmax), 224 ±= 29 W) were recruited to participate. Participants were fully informed of the study procedures and their right to withdraw before providing written consent to participate. The study was pre-approved by the National Health Service Research Ethics Committee, Black Country, West Midlands, UK (17/WM/0321) and was conducted in accordance with the Declaration of Helsinki.

### Experimental overview

Participants attended the laboratory on five occasions. Prior to the experimental periods, participants attended the laboratory for a pre-testing visit to determine VO_2_peak and Wmax. The experimental period then consisted of two identical experimental blocks in which participants visited the laboratory before and after a seven-day supplementation period. During the supplementation period participants received either 1000 mg·d^-1^ nicotinamide riboside (NR; Niagen, ChromaDex, Irvine CA, USA) or 1000 mg·d^-1^ of a cellulose placebo (PLA) in a double-blinded, randomised, counter-balanced, crossover design. Supplements were consumed twice daily such that participants were instructed to consume 500 mg of supplement at ∼9 am and ∼9 pm each day. A two-week washout period was employed between experimental blocks. After measuring height (Seca 220, Seca, Birmingham, UK) and body mass (Champ II, OHAUS, Griefensee, Switzerland) participants performed a graded exercise test to exhaustion on a cycle ergometer (Lode Excalibur, Groningen, Netherlands). The test began at 50 W with power increasing by 25 W every three minutes thereafter. Respiratory variables were measured continuously during exercise using a breath-by-breath metabolic cart (Vyntus CPX, Jaeger, CareFusion, Germany), heart rate was monitored throughout (RCX5, Polar Electro Oy, Kempele, Finland) and ratings of perceived exertion (RPE) were determined using a 6-20 Borg scale during the final 15 seconds of each 3-minute stage [23]. VO_2_peak was determined as the highest rolling 30-second average of VO_2_. Wmax was determined as work rate at the last completed stage plus the fraction of time spent in the final non-completed stage multiplied by the increment in work rate (25W).

### Experimental trials

Participants refrained from alcohol for 72 h, caffeine for 24 h and exercise for 48 h prior to each experimental trial. For 72 h prior to each experimental trial participants consumed a replicated diet. For the first 48 h of this period participants consumed a diet that replicated their ad libitum intake recorded via a weighed food diary prior to the first experimental visit. For the final 24 h prior to each experimental visit participants were provided with a standardised fixed energy intake diet (energy: 2271 kcal; macronutrient composition: 63% carbohydrate, 21% fat and 16% protein). For the pre-supplementation visit, participants arrived at the laboratory at ∼8:30am following an ∼12-hour overnight fast. Upon arrival, participants rested in the supine position for approximately five minutes before a venous blood sample was collected via venepuncture from an antecubital forearm vein. A resting skeletal muscle biopsy was then taken from the medial vastus lateralis. Participants then consumed the first 500 mg dose of their supplement prior to leaving the laboratory. For the post-supplementation visit, participants arrived at the laboratory at ∼7:30 am following an ∼12-hour overnight fast. Participants rested in the supine position for ten minutes prior to a 20-minute measurement of resting metabolic rate under a ventilated hood using the GEMNutrition indirect calorimeter (GEMNutrition, Daresbury, UK). A cannula was then inserted into an antecubital forearm vein and a baseline venous blood sample was collected prior to providing a pre-exercise skeletal muscle biopsy from the medial vastus lateralis.

Participants then cycled for one-hour at 60% Wmax before a second skeletal muscle biopsy was taken immediately post-exercise (completed within two minutes of exercise cessation) after which they rested in a supine position prior to a third skeletal muscle biopsy obtained three-hours post-exercise. A new incision was made for each biopsy at least 2 cm proximal from the previous site. Venous blood was collected throughout rest periods and during exercise. Respiratory variables were measured pre-exercise and at 15-minute intervals throughout exercise, heart rate was monitored continuously throughout exercise and RPE was determined at 15-minute intervals throughout exercise. Carbohydrate and fat oxidation were calculated from VO_2_ and VCO_2_ using the moderate-high exercise intensities equation of Jeukendrup and Wallis [24] during exercise and Frayn [25] at rest. Participants were allowed to drink water ad libitum during rest and exercise periods during the visit following the first supplementation period, which was matched during the second experimental trial.

### Muscle biopsies

Muscle biopsies were obtained from separate incision sites on the medial vastus lateralis under local anaesthesia (1% lidocaine; B. Braun, Melsungen, Germany) by a Bergström needle adapted with suction. Muscle was rapidly blotted to remove excess blood and was immediately flash frozen in liquid nitrogen. In the case of pre-supplementation and pre exercise biopsies, an ∼20mg section was removed prior to freezing and placed in ice-cold BIOPS buffer (2.77 mM CaK2EGTA, 7.23 mM K2EGTA, 5.77 mM Na2ATP, 6.56 mM MgCl2, 20 mM taurine, 15 mM Na2Phosphocreatine, 20 mM imidazole, 0.5 mM DTT, and 50 mM MES) for the immediate measurement of mitochondrial respiration. Frozen muscle was powdered using a Cellcrusher tissue pulverizer on dry ice and stored at −80°C prior to analysis.

### High-resolution respirometry

Skeletal muscle fibres were mechanically separated under a light microscope and permeabilised by incubation in BIOPS buffer containing 50 mg·ml^-1^ of saponin for 30 minutes followed by a 15-minute wash in MiR05 buffer (0.5 mM EGTA, 3 mM MgCl2.6H2O, 60 mM K lactobionate, 20 mM taurine, 10 mM KH2PO4, 20 mM HEPES, 110 mM sucrose, and 1 g.L-1 fatty acid-free bovine serum albumin). Samples were then weighed and analysed in duplicate using an Oroboros O2K (Oroboros Instruments, Innsbruck, Austria). When substantial variability was apparent between duplicates a third sample was run. Data was collected at 37°C in hyperoxygenated (200-400 μM) conditions in MiR05 buffer. The substrate-uncoupler inhibitor titration performed was as follows: 5 mM pyruvate, 2 mM malate, and 10 mM glutamate was added to measure leak respiration through complex one (CIL); 5 mM ADP was then added to measure coupled oxidative phosphorylation through complex one (CIP); 10 mM succinate was then added to measure coupled oxidative phosphorylation through complexes one and two (CI+IIP); 10 μM cytochrome-c was added to test outer mitochondrial membrane integrity; titrations of 0.5 μM FCCP until maximal respiration were then added to measure maximal electron transport chain capacity (CI+IIE); 5 μM antimycin A was then added to measure nonmitochondrial respiration. Respiration was normalised to tissue masses and non-mitochondrial respiration was subtracted to give mass-specific mitochondrial respiration. In all samples the increase in respiration following addition of cytochrome-c was less than 10%, indicating preserved mitochondrial membrane integrity.

### Immunoblotting

Tissue was homogenized in a 10-fold mass excess of ice-cold sucrose lysis buffer (50 mM tris, 1 mM EDTA, 1 mM EGTA, 50 mM NaF, 5 mM Na4P2O7-10H_2_O, 270 mM sucrose, 1 M triton-X, 25 mM β-glycerophosphate, 1 μM trichostatin A, 10 mM nicatinamide, 1mM 1,4 dithiothreitol, 1% phosphatase inhibitor Cocktail 2; Sigma, 1% phosphatase inhibitor cocktail 2; Sigma, 4.8% cOmplete mini protease inhibitor cocktail; Roche) using an IKA T10 basic ULTRA-TURRAX homogeniser (IKA, Oxford, UK) followed by shaking at 4°C for 30 minutes and centrifuging at 4°C and 8000 g for 10 minutes to remove insoluble material. Protein concentrations were determined by the DC protein assay (Bio-Rad, Hercules, California, USA). Samples were prepared in laemmli sample buffer, boiled at 97°C for 5 min (with the exception of an aliquot set aside for determination of electron transport chain protein content which remained unboiled) and an equal volume of protein (18-36 μg) was separated by SDS-PAGE on 8 - 12.5% gels at a constant current of 23 mA per gel. Proteins were transferred on to BioTrace NT nitrocellulose membranes (Pall Life Sciences, Pensacola, Florida, USA) via wet transfer at 100 V for one hour. Membranes were then stained with Ponceau S (Sigma-Aldrich, Gillingham, UK) and imaged to check for even loading. Membranes were blocked in 3% dry-milk in tris-buffered saline with tween (TBST) for one hour before being incubated in primary antibody overnight at 4°C. Membranes were washed in TBST three times prior to incubation in appropriate horse radish peroxidase (HRP)-conjugated secondary antibody at room temperature for one hour. Membranes were then washed in TBST three times prior to antibody detection via enhanced chemiluminescence HRP substrate detection kit (Millipore, Watford, UK). Imaging and band quantification were undertaken using a G:Box Chemi-XR5 (Syngene, Cambridge, UK).

### Antibodies

All primary antibodies were used at a concentration of 1:1000 in TBST unless otherwise stated. Pan-acetylation (ab193), ac-MnSOD^K122^ (ab214675) and OXPHOS cocktail (ab110411) were purchased from abcam; AMP-activated protein kinase alpha (AMPKα; 2603), p-AMPK^Thr172^ (2535), p-ACC^Ser79^ (3661), calmodulin dependent kinase II (CAMKII; 3362), p-CAMKII^Thr268^ (12716), cAMP response element binding protein (CREB; 1°: 1:500; 9197), p-CREB^Ser133^ (1°: 1:500; 9191), glyceraldehyde 3-phosphate dehydrogenase (GAPDH; 1:5000; 2118), p38 mitogen activated protein kinase (p38 MAPK; 9212), p-p38 MAPK^Thr180/Tyr182^ (4511), poly ADP-ribose polymerase 1 (PARP1; 1°: 1:500; 9542), tumour protein 53 (p53; 2°: 1:2000; 2527) and acp53^K382^ (1°: 1:500 in 3% BSA, 2°: 1:2000; 2570) were purchased from Cell Signaling Technology; acetyl CoA carboxylase (ACC; 05-1098), superoxide dismutase (MnSOD; 1°: 1:2000; 06-984), PGC-1α (ab3242) and poly-ADPribose (PAR; 1°: 1:500; MABE1031) were purchased from Merck Millipore. Secondary antibodies were used at a concentration of 1:10000 in TBST unless otherwise stated. Anti-rabbit (7074) and anti-mouse (7076) antibodies were from Cell Signaling Technology.

### Real time RT-qPCR

RNA was extracted from ∼20 mg of muscle by homogenising in 1 mL of Tri reagent (Sigma Aldrich, Gillingham, UK) using an IKA T10 basic ULTRATURRAX homogeniser (IKA, Oxford, UK). Phase separation was achieved by addition of 200 μL of chloroform and centrifugation at 12000 g for 15 minutes. The RNA-containing supernatant was removed and mixed with an equal volume of 2-propanol. RNA was purified on Reliaprep spin columns (Promega, Madison, Wisconsin, USA) using the manufacturer’s instructions, which includes a DNase treatment step. RNA concentrations were determined using the LVis function of the FLUOstar Omega microplate reader (BMG Labtech, Aylesbury, UK). RNA was diluted to 30 ng·μL^-1^ and reverse transcribed to cDNA in 20 μL volumes using the nanoScript 2 RT kit and oligo(dT) primers (Primerdesign, Southampton, UK) as per the manufacturer’s instructions. RT-qPCR analysis of mRNA content was performed in triplicate by using Primerdesign custom designed primers (Table 1) and commercially available ACTB, B2M GAPDH, (Primerdesign) and Precision plus qPCR Mastermix with low ROX and SYBR (Primerdesign) on a QuantStudio3 Real-Time PCR System (Applied Biosystems, Thermo Fisher, UK). The qPCR reaction was run as per the manufacturer’s instructions (Primerdesign) and followed by a melt curve (Applied Biosystems) to ascertain specificity. 2-20 ng of cDNA was added to each well in a 20 μL reaction volume. qPCR results were analysed using Experiment Manager (Thermo Fisher). mRNA expression is expressed relative to the expression in the pre-exercise sample during FED for each individual using the 2-ΔΔCQ method [26] with the geometric mean of Cq values for ACTB, B2M and GAPDH used as an internal control [27]. Optimal stability of housekeeper genes was determined using RefFinder [28]. Statistical analyses were performed on log-transformed ΔΔCQ values.

**Table 1.**
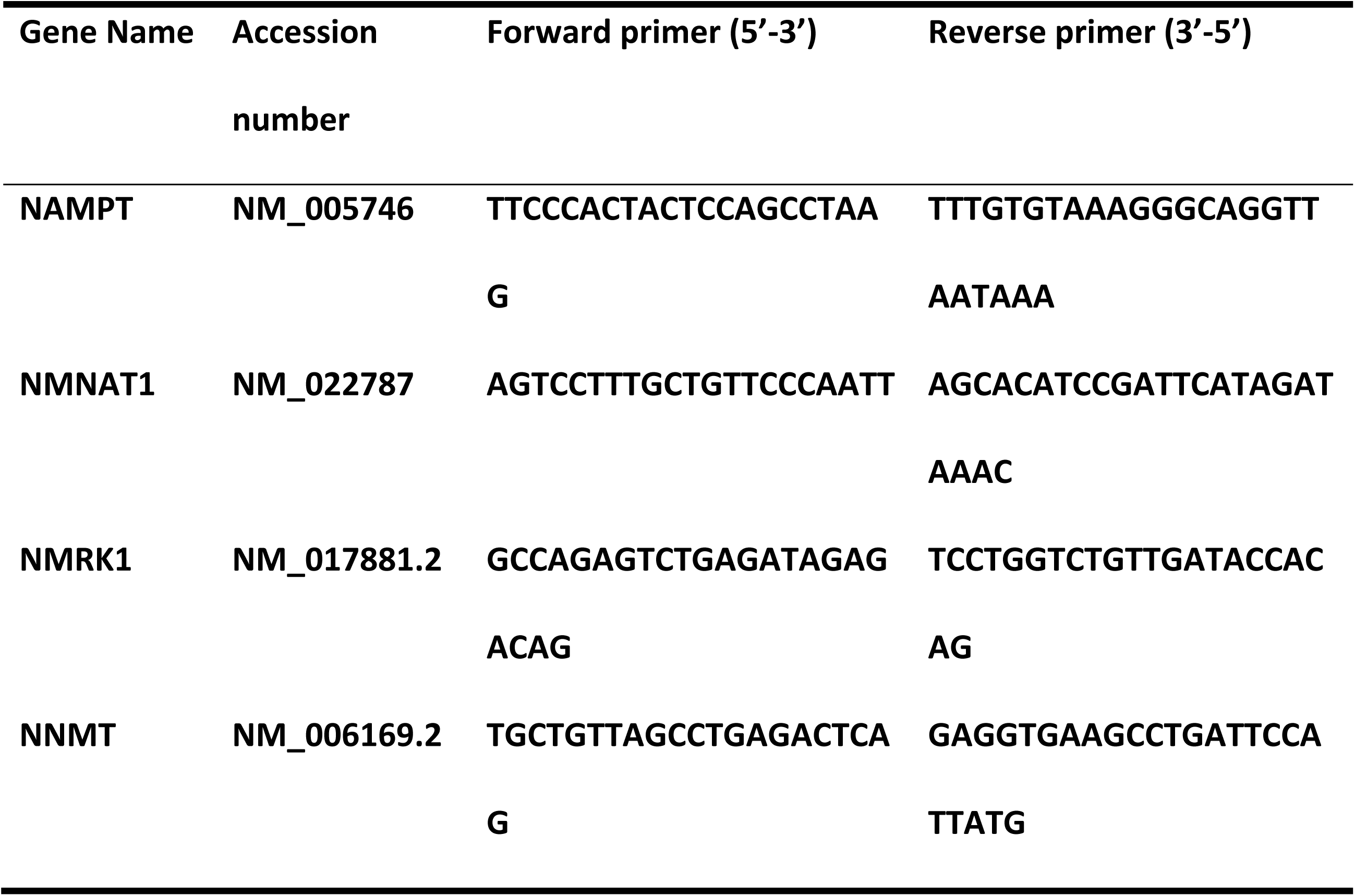
Gene Accession numbers and corresponding primer sequences.

### Blood analyses

Blood samples were collected into tubes containing ethylenediaminetetraacetic acid (EDTA; BD, Oxford, UK) for the collection of plasma. Samples were placed immediately upon ice prior to centrifugation at 1600 g at 4°C for 10 minutes before collection of plasma from the supernatant. Plasma was frozen at −80°C until further analysis. Plasma samples were subsequently analysed on an autoanalyser (iLAB650, Instrumentation Laboratory, Bedford, MA, USA) for glucose, lactate, non-esterified fatty acid (NEFA) and glycerol (Randox Laboratories, County Antrim, UK) using commercially available kits.

### Statistics

Two-way repeated measures ANOVAs assessed effects of time, treatment and time*treatment interaction effects for all time-course data. Ryan-Holm-Bonferroni multiple comparison corrections were applied post-hoc where applicable. Differences in means for resting and exercising VO_2_, VCO_2_, respiratory exchange ratio (RER), substrate utilisation, heart rate and RPE were assessed using repeated-measures t-tests. All statistics were performed using the Statistical Package for the Social Sciences (SPSS) version 22.0. Data are presented as means with 95% confidence intervals. Statistical significance was accepted as p <0.05.

## Results

### Substrate utilisation and systemic availability

Seven days of NR supplementation did not influence resting metabolic rate (PLA: 1859 ± 202 vs NR: 1772 ± 211 kcal·d^-1^; p = 0.486). Furthermore, substrate utilisation at rest was similar following supplementation of NR or PLA (carbohydrate oxidation: PLA: 0.09 ± 0.04 vs NR: 0.11 ± 0.03 g·min^-1^; p = 0.446, fat oxidation: PLA: 0.10 ± 0.03 vs NR: 0.09 ± 0.02 g·min^-1^; p = 0.395, RER: PLA: 0.79 ± 0.04 vs NR: 0.80 ± 0.03; p = 0.563). Carbohydrate and fat oxidation during exercise were also similar between trials (Table 2). VO_2_, VCO_2_, RER, heart rate and RPE did not differ between trials during exercise (Table 2). There was no effect of NR on resting or exercising plasma NEFA, glycerol, glucose or lactate (Figure 1). Plasma NEFA concentration initially decreased during the first 15 minutes of exercise before returning to pre-exercise values for the remainder of the exercise bout (main effect of treatment; p = 0.891, time; p < 0.001, interaction; p = 0.296). Following exercise (80 minutes) plasma NEFA concentration increased and remained elevated above pre-exercise values from 120 minutes until the end of the trial (240 minutes). Plasma glycerol concentration increased during exercise and remained elevated above pre-exercise values for the remainder of the trial (main effect of treatment; p = 0.106, time; p < 0.001, interaction; p = 0.720). Plasma glucose was marginally, although significantly, decreased from pre-exercise values at two hours after the cessation of exercise (main effect of treatment; p = 0.175, time; p = 0.010, interaction; p = 0.174). Plasma lactate increased during exercise and remained elevated for the first 20 minutes of recovery (main effect of treatment; p = 0.192, time; p = 0.001, interaction; p = 0.585).

**Table 2.**
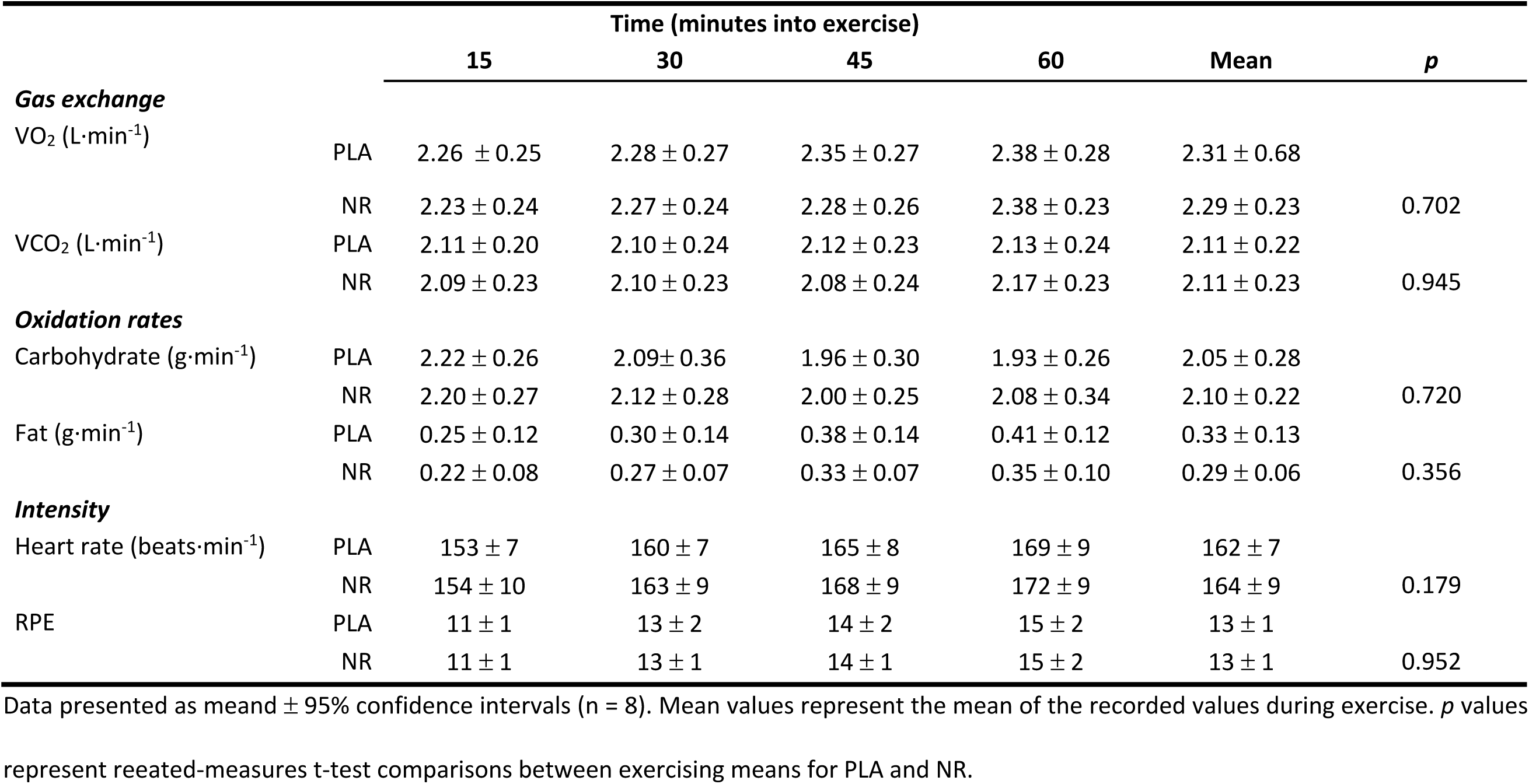
Effect of NR supplementation on cardio-respiratory changes and substrate utilisation during exercise.

**Figure 1.**
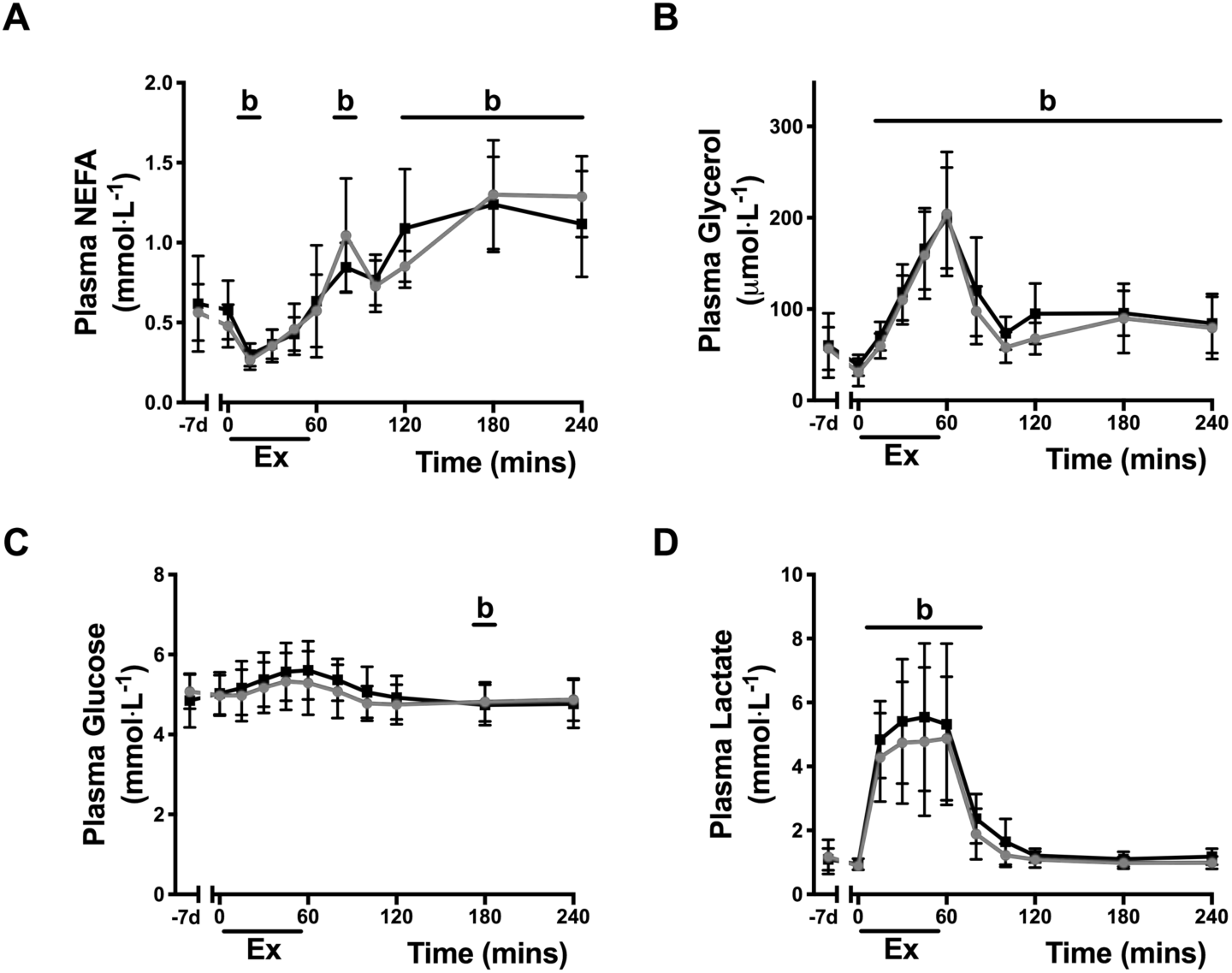
NR supplementation does not alter plasma NEFA, glycerol, glucose or lactate at rest or during exercise. Time-course for plasma NEFA (A.), glycerol (B.), glucose (C.) and lactate (D.) in PLA (black) and NR (grey). b: main effect of time (significantly different to pre-exercise; p ≤ 0.05. Data presented as means ± 95% confidence intervals (n = 8).

### Skeletal muscle mitochondrial function and protein content

Rates of mitochondrial respiration were similar to those previously reported [29, 30]. There were no changes observed in CIL (main effect of treatment; p = 0.319, time; p = 0.833, interaction; p = 0.588), CIP (main effect of treatment; p = 0.979, time; p = 0.388, interaction; p = 0.551), CI+IIP (main effect of treatment; p = 0.612, time; p = 0.216, interaction; p = 0.993) or CI+IIE (main effect of treatment; p = 0.657, time; p = 0.190, interaction; p = 0.621) respiration following supplementation of NR or PLA (Figure 2A). Furthermore, the content of proteins within each of the five electron transport chain complexes were unchanged following NR or PLA supplementation (Figure 2B; p > 0.05).

**Figure 2.**
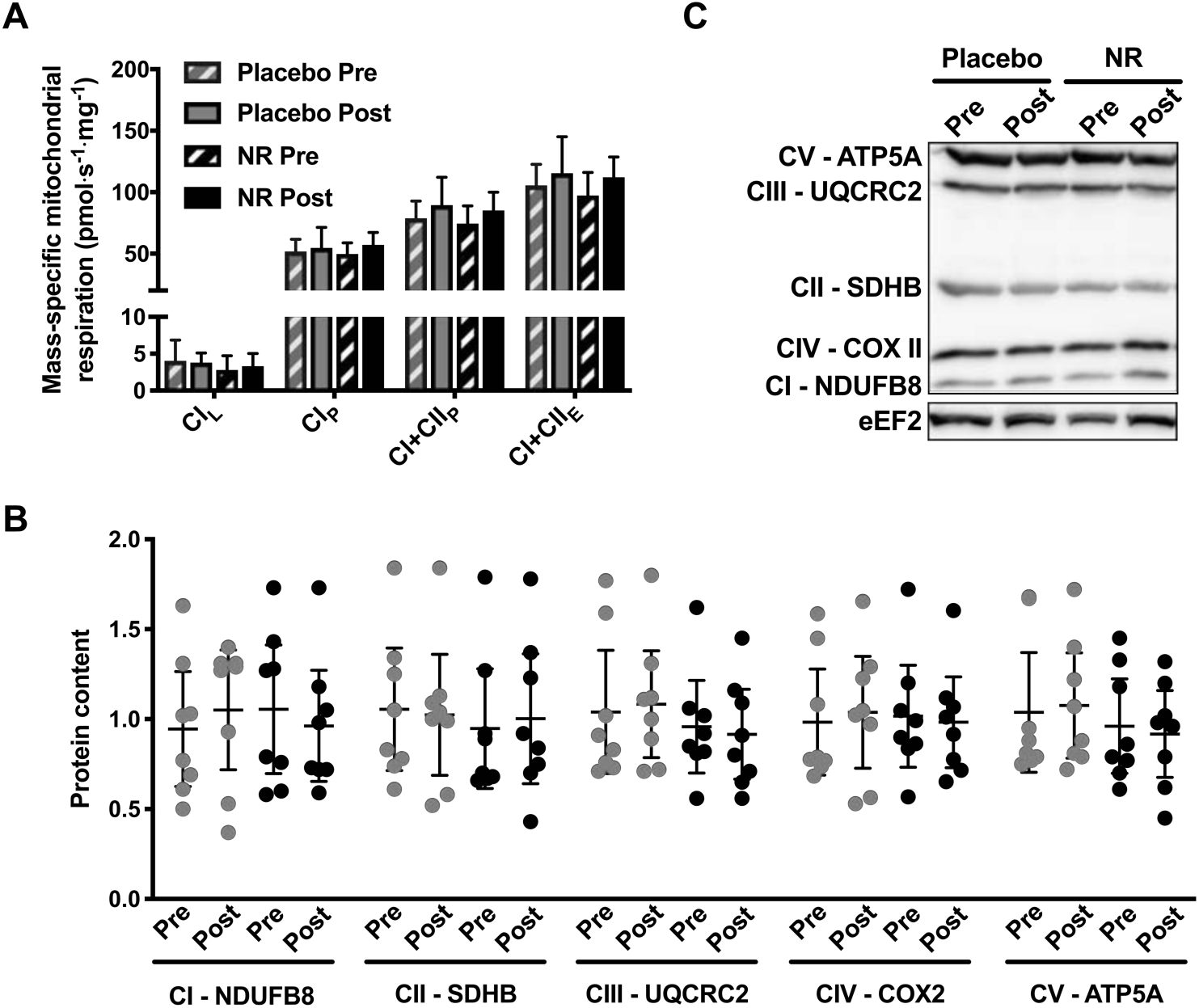
Seven days of NR supplementation does not induce mitochondrial biogenesis in skeletal muscle. A. There were no changes in the mass-specific mitochondrial leak respiration through complex I (CIL), coupled respiration through complex I (CIP), coupled respiration through complexes I and II (CI+IIP), or maximal electron transport chain capacity (CI+IIE) following seven days of NR supplementation (p > 0.05). B. Similar content of proteins within the five electron transport chain complexes pre- and post-supplementation of PLA (grey) or NR (black) (p > 0.05). C. Representative immunoblot images. Data presented as means ± 95% confidence intervals (n = 8).

### Skeletal muscle signalling

Global acetylation within skeletal muscle was unaffected by NR supplementation or exercise (Figure 3A; main effect of treatment; p = 0.845, time; p = 0.120, interaction; p = 0.106). Furthermore, the acetylation of p53^Lys382^, a SIRT1 deacetylation target [31], and MnSOD^K122^, a SIRT3 deacetylation target [32], were unchanged throughout the intervention (Figure 3C & D; p53^Lys382^: main effect of treatment; p = 0.723, time; p = 0.786, interaction; p = 0.354, MnSOD^K122^: main effect of treatment; p = 0.324, time; p = 0.409, interaction; p = 0.332). The protein content of PARP1 was unaffected by NR supplementation as post-hoc analyses revealed no significant difference despite a significant treatment*time interaction effect (main effect of treatment; p = 0.498, time; p = 0.520, interaction; p = 0.040; Figure 4A). Auto-PARylation of PARP1, a proxy of PARP1 activity [33], was also unchanged by NR or exercise (main effect of treatment; p = 0.512, time; p = 0.255, interaction; p = 0.115; Figure 4B).

**Figure 3.**
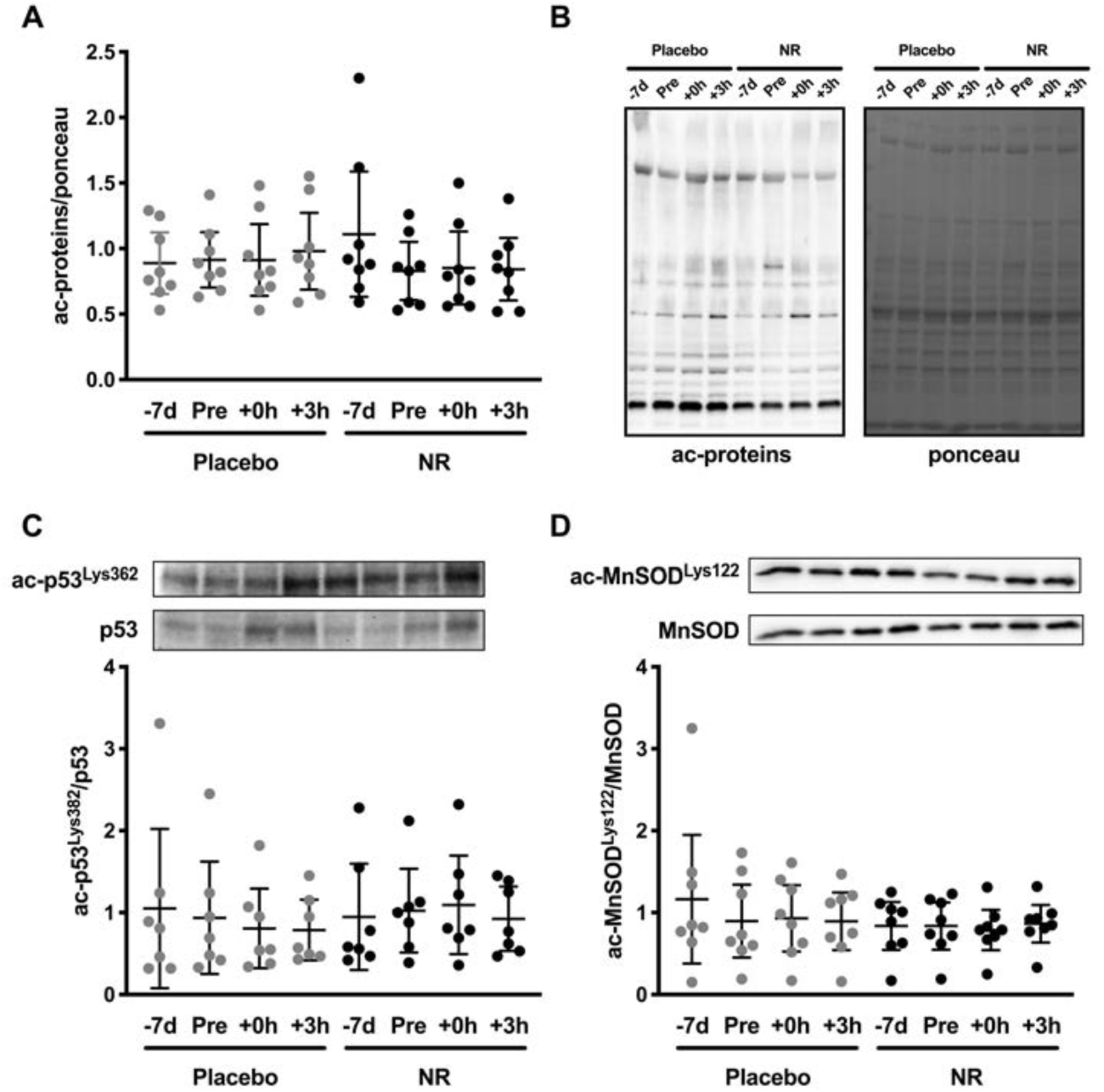
Seven days NR supplementation does not influence Sirtuin deacetylase activity at rest or following endurance exercise. A. Global acetylation within skeletal muscle is unaffected by NR supplementation or exercise (n = 8; p > 0.05). B. Representative immunoblot images of global acetylation and Ponceau S stain. C. Acetylation of p53^Lys382^, a SIRT1 deacetylation site, is unchanged by NR supplementation at rest or following endurance exercise (n = 7; p > 0.05). D. Acetylation of MnSOD^Lys122^, a SIRT3 deacetylation site, is unchanged by NR supplementation at rest or following endurance exercise (n = 8; p > 0.05). - 7d: pre-supplementation; Pre: pre-exercise (post-supplementation); +0h: immediately post-exercise; +3h: three hours post-exercise. All values are presented relative to the group mean for all pre-supplementation samples. Data presented as means ± 95% confidence intervals.

**Figure 4.**
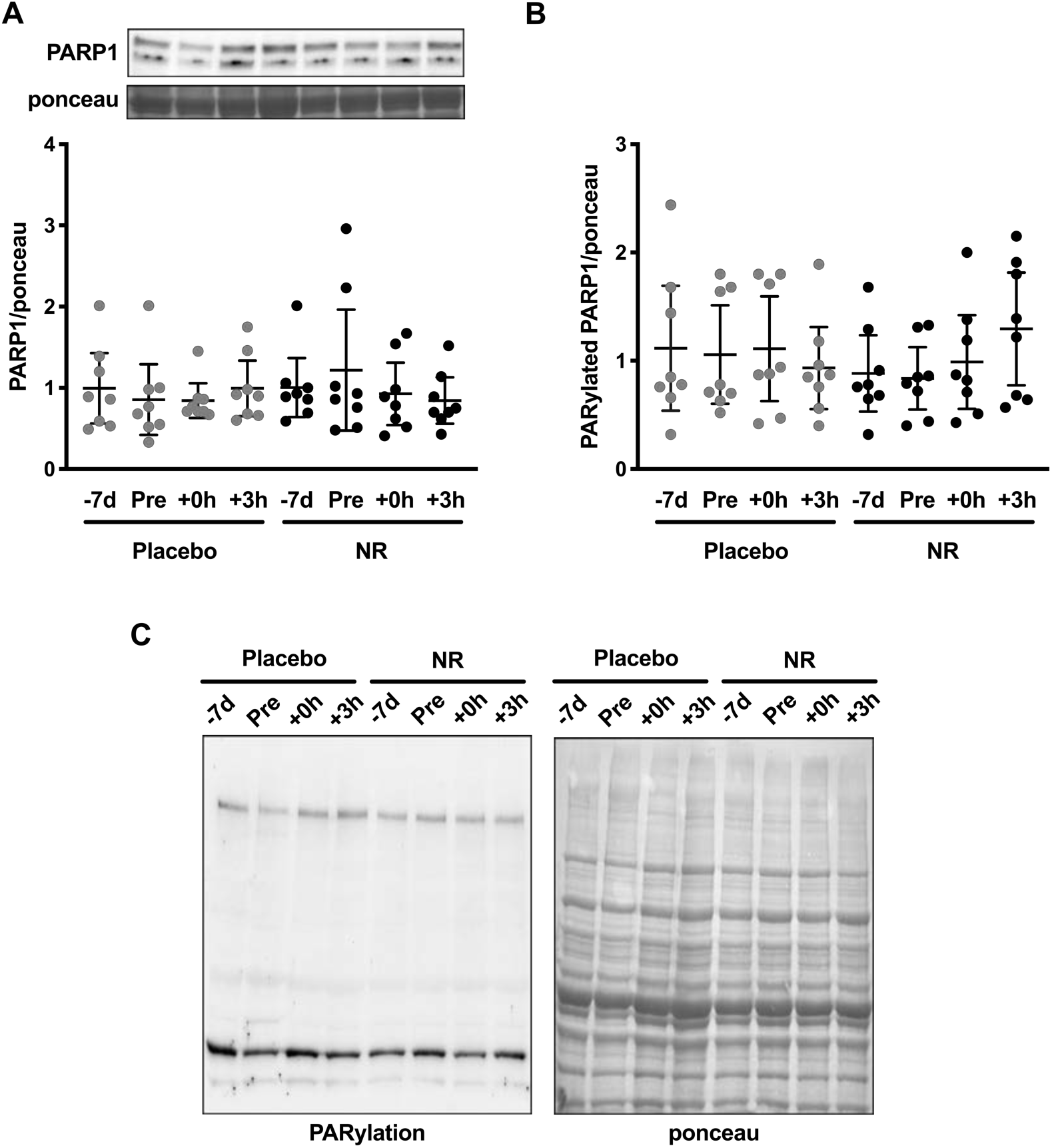
Seven days of NR supplementation does not influence PARP1 protein content or PARylated PARP1 protein content. PARP1 protein content (A.) and auto-PARylation of PARP1 (B.) are unaffected by NR supplementation or exercise (p < 0.05). C. Representative immunoblot images of PARylation and Ponceau S stain. -7d: pre-supplementation; Pre: preexercise; +0h: immediately post-exercise; +3h: three hours post-exercise. All values are presented relative to the group mean for all pre-supplementation samples. Data presented as means ± 95% confidence intervals (n = 8).

Exercise increased the phosphorylation of AMPK^Thr172^ (Figure 5A, main effect of time; p = 0.002) by ∼1.6-fold immediately post-exercise (p = 0.031 vs pre-exercise). There was no effect of treatment (p = 0.216) or a treatment*time interaction effect (p = 0.472). Phosphorylation of ACC^Ser79^ (Figure 5B) increased ∼4-fold immediately post-exercise (p < 0.001 vs pre-exercise) and remained ∼1.4-fold elevated 3-h post-exercise (p = 0.013 vs pre-exercise, main effect of time; p < 0.001). CREB^Ser133^ phosphorylation was unaffected by exercise or NR (main effect of treatment; p = 0.651, time; p = 0.462, interaction; p = 0.810; Figure 5C). p38 MAPK^Thr180/Tyr182^ phosphorylation was not significantly affected by exercise or NR (Figure 5D), as post-hoc analyses revealed no significant differences despite a treatment*time interaction effect (main effect of treatment; p = 0.124, time; p = 0.942, interaction; p = 0.034). CAMKII^Thr286^ phosphorylation was not altered by exercise or NR (main effect of treatment; p = 0.574, time; p = 0.177, interaction; p = 0.236; Figure 5E).

**Figure 5.**
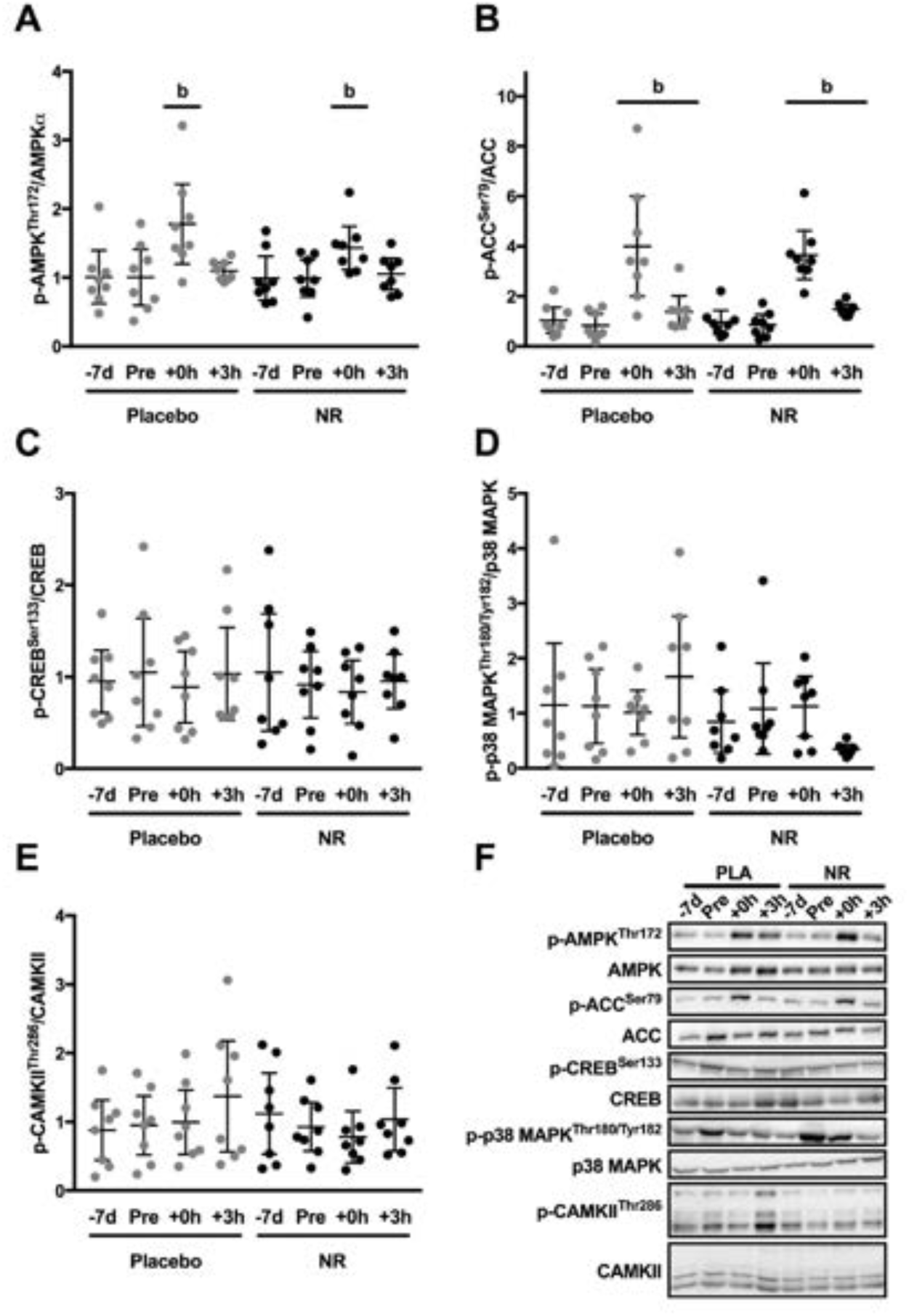
Activation of exercise-sensitive signalling pathways following NR supplementation and endurance exercise. A. Phosphorylation of AMPK^Thr172^ is increased immediately post exercise in each trial. B. Phosphorylation of ACC^Ser79^ is increased immediately postexercise and remains elevated three hours post-exercise in each trial. C. CREB^Ser133^, D. p38 MAPK^Thr180/Tyr182^ and E. CAMKII^Thr286^ remain unchanged throughout the intervention. F. Representative immunoblot images. -7d: presupplementation; Pre: pre-exercise (post supplementation); +0h: immediately post-exercise; +3h: three hours post-exercise. b: main effect of time (significantly different to pre-exercise; p ≤ 0.05). All values are presented relative to the group mean for all pre-supplementation samples. Data presented as means ± 95% confidence intervals (n = 8).

### Metabolic mRNA response

Seven days of NR supplementation did not alter resting PPARGC1A mRNA expression in skeletal muscle (Figure 6A). PPARGC1A mRNA increased ∼5-fold three hours post-exercise (p = 0.025 vs pre-exercise, main effect of time; p = 0.003). Post-exercise PPARGC1A mRNA expression was similar in PLA and NR trials (main effect of treatment; p = 0.257, interaction; p = 0.591). Expression of pyruvate dehydrogenase kinase 4 (PDK4; Figure 6B) increased post-exercise (main effect of time; p = 0.001) and was ∼10-fold elevated three hours post-exercise (p = 0.029 vs pre-exercise). mRNA expression of PDK4 was similar between PLA and NR trials (main effect of treatment; p = 0.827, interaction; p = 0.521).

**Figure 6.**
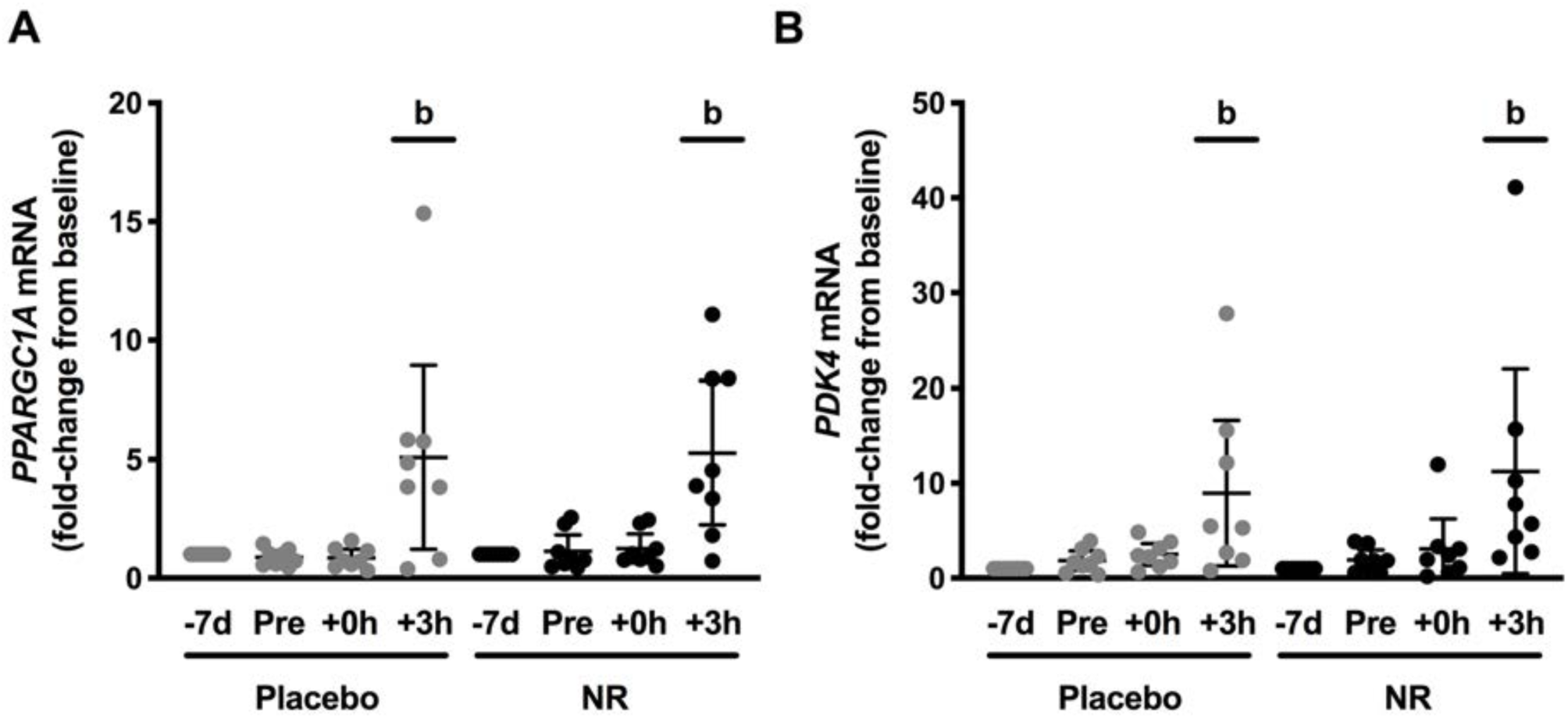
Seven days of NR supplementation does not alter resting or exercise-induced PGC-1α or PDK4 mRNA expression. A. Resting and exercise induced PGC-1α mRNA expression is similar between NR and PLA trials. B. Resting and exercise induced PDK4 mRNA expression is similar between NR and PLA trials. -7d: pre-supplementation; Pre: pre-exercise (post-supplementation); +0h: immediately post-exercise; +3h: three hours post-exercise. b: main effect of time (significantly different to pre exercise; p ≤ 0.05). All values are presented relative to individual pre supplementation values for each trial and reported as means ± 95% confidence intervals (n = 8).

### mRNA expression of enzymes within the NAD^+^ synthesis and salvage pathways

NR supplementation did not alter the mRNA expression of nicotinamide riboside kinase 1 (NMRK1; main effect of treatment; p = 0.432) within skeletal muscle (Figure 7A). NMRK1 mRNA expression did show a tendency for a main effect of time (p = 0.071). There was no treatment*time interaction effect for NMRK1 mRNA (p = 0.203). mRNA expression of NAMPT, the rate limiting enzyme in NAD^+^-salvage (8, 12, 14), was unaffected by NR supplementation or exercise (Figure 7B; main effect of treatment; p = 0.303, time; p = 0.305, interaction; p = 0.442). Nicotinamide mononucleotide acetyl transferase 1 (NMNAT1) mRNA expression was not influenced by NR supplementation (Figure 7C), however showed a trend to decrease three hours post-exercise (p = 0.065 vs preexercise, main effect of time: p = 0.046). There was no effect of treatment (p = 0.482) nor a treatment*time interaction effect (p = 0.168). Nicotinamide N-methyltransferase (NNMT) increased in expression three-hours post-exercise (Figure 7D; p = 0.010 vs pre-exercise, main effect of time: p = 0.001). However, the post-exercise mRNA expression of NNMT was suppressed following NR supplementation (treatment*time interaction: p = 0.029), such that the exercise-induced NNMT mRNA expression was only increased in the PLA trial (PLA 3h post-exercise vs PLA pre-exercise: p = 0.010), while there was also a trend towards a difference between NR and PLA three hours post-exercise (p = 0.116). There was no main effect of treatment (p = 0.148).

**Figure 7.**
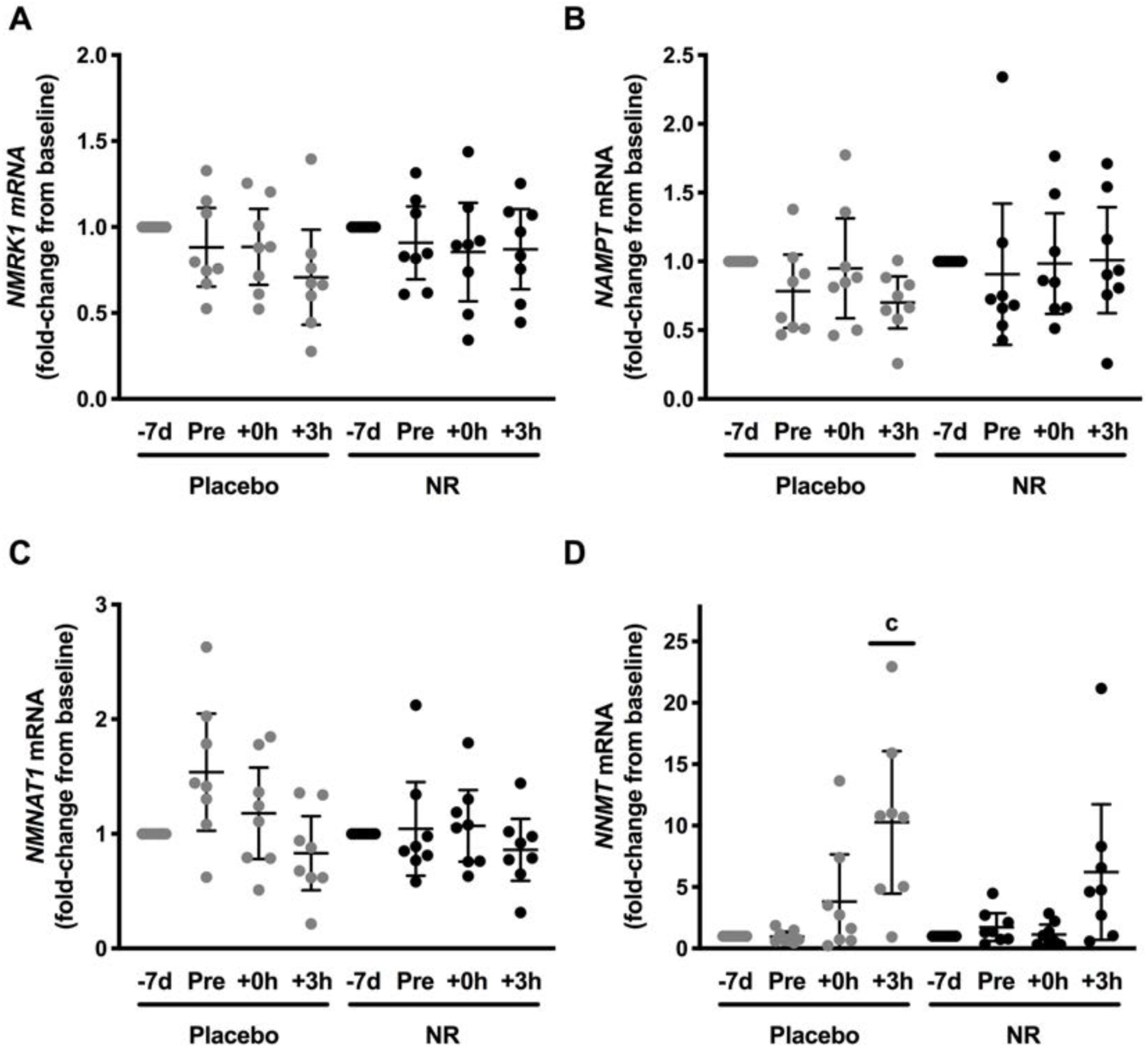
mRNA expression of enzymes in the NAD+ synthesis and salvage pathways within skeletal muscle following NR supplementation and endurance exercise. mRNA expression of A. NMRK1 and B. NAMPT were unaffected by NR supplementation or endurance exercise. C. NMNAT1 mRNA expression displayed a tendency to decrease three hours post-exercise (p = 0.065). D. mRNA expression of NNMT increased three hours post-exercise in PLA but this was impaired following NR supplementation. -7d: pre-supplementation; Pre: pre-exercise (post supplementation); +0h: immediately post-exercise; +3h: three hours post-exercise. c: interaction effect (different to pre-exercise within Nutrition and post-exercise energy-sensing in skeletal muscle treatment; p ≤ 0.05). All values are presented relative to individual pre supplementation values for each trial and reported as means ± 95% confidence intervals (n = 8).

## Discussion

Contrary to our hypothesis, the activity of the NAD^+^-dependent deacetylases SIRT1 and SIRT3 and the mRNA expression of PPARGC1A were unaffected by NR supplementation, both at rest and in the post-exercise recovery period. Furthermore, seven days of NR supplementation (1000 mg·d^-1^) did not alter whole-body metabolism or substrate utilisation. Finally, and somewhat surprisingly, NR impaired the exercise-induced increase of NNMT mRNA expression, an enzyme putatively involved in regulating whole-body fatty acid metabolism [34].

Previous work in rodents supplemented with NR for periods of 4-16 weeks have reported an increase in skeletal muscle NAD^+^ content in parallel to increased SIRT1 and SIRT3 activity and mitochondrial biogenesis [7, 8, 12, 13]. However, in the current study, NR supplementation did not alter sirtuin activity or PPARGC1A mRNA expression in human skeletal muscle at rest or following endurance exercise. NAD^+^-dependent signalling does appear to be more tightly regulated in human compared to rodent skeletal muscle [16]. For example, endurance exercise increases SIRT1 activity in mice (assessed via p53 deacetylation) [35] but does not appear to produce the same response in humans (Figure 3), possibly indicative of slower NAD^+^ turnover in human skeletal muscle or additional levels of regulation [16]. In addition, the protein content and activity of the NAD^+^-dependent protein PARP1 was also unchanged by exercise or NR supplementation in the current study. However, this does support rodent data showing that NR supplementation does not alter basal PARylation within skeletal muscle [7, 8]. Therefore, NR supplementation appears less efficient in altering skeletal muscle NAD^+^-content in healthy human skeletal muscle [18, 20, 21] when compared to rodent skeletal muscle, where effects of supplementation are more pronounced (∼10% increase in NAD^+^) [7] and even more so when NR is provided during metabolic stress (∼30% increase in NAD^+^) [7]. We have previously shown that 21 days NR supplementation increases MeNAM and NAAD content in old human skeletal muscle indicative of altered NAD^+^ flux [20, 21], however did not observe changes in NAD^+^ content or mitochondrial respiration [20, 21]. Collectively, therefore, NAD^+^ metabolism in human skeletal muscle seems to be more tightly regulated and resistant to precursor supplementation than in rodents.

NR supplementation for one week did not alter circulating substrate availability or whole-body substrate utilisation either at rest or during 60% Wmax cycling in healthy recreationally active males. This is in accordance with recent reports where NR supplementation of 1000 mg·d^-1^ for six weeks or 2000 mg·d^-1^ for 12 weeks had no effect on resting energy expenditure, substrate utilisation or fasting concentrations of glucose or NEFA [6, 17, 20, 21]. Furthermore, six weeks of NR supplementation does not alter RER during an incremental exercise test in elderly males [6]. However, these data are in contrast to rodent studies, which have demonstrated that NR supplementation can increase metabolic flexibility [10] and fat oxidation during the inactive phase [7], which occurs alongside induced mitochondrial biogenesis [7]. Changes in substrate utilisation with NR supplementation may therefore be a physiological outcome of mitochondrial biogenesis. Indeed, in the current study, no changes in skeletal muscle mitochondrial respiration or content of electron transport chain proteins were apparent, which perhaps is unsurprising given the relatively short supplementation period. However, 6 and 12 weeks of NR supplementation also failed to increase skeletal muscle mitochondrial volume or respiratory capacity in humans [18, 20, 21]. In contrast, two weeks of acipimox administration (750 mg·d^-1^), a nicotinic acid-derivative, increased skeletal muscle mitochondrial respiratory capacity in type II diabetes [36], whilst four months of nicotinic acid supplementation increased mitochondrial mass and cytochrome c oxidase activity in healthy participants and patients with mitochondrial myopathies [37]. It remains unclear why nicotinic acid may be more potent than NR in stimulating mitochondrial biogenesis in human skeletal muscle.

Our data indicates that exercise and NR alter the NAD^+^-consumption/salvage machinery within skeletal muscle. The mRNA expression of NNMT, a methyltransferase of nicotinamide (NAM) that produces methylated NAM (MeNAM) and prevents NAD^+^-salvage [38], is increased following endurance exercise although this response was impaired by NR supplementation. Previous studies have also shown an increase in skeletal muscle NNMT mRNA and/or protein expression following endurance exercise training in rats [39] and four days of energy restriction in humans [34]. Ström et al [34] went on to demonstrate elevated skeletal muscle NNMT mRNA expression coincided with an increase in circulating MeNAM. In addition, plasma MeNAM concentrations are increased following a single bout of endurance exercise in mice, an effect that could only be partially explained by increased NNMT activity in the liver [40]. MeNAM can be secreted from human primary myotubes and can induce lipolysis in rat primary adipocytes [34]. However, despite elevations in systemic and skeletal muscle MeNAM during NR supplementation in humans [20-22], whole-body fatty acid availability at rest, during exercise and during the post-exercise recovery period are unaffected by NR supplementation. The reduction in exercise-induced skeletal muscle NNMT mRNA expression following NR supplementation is a particularly surprising finding given elevated plasma and skeletal muscle MeNAM concentrations during NR supplementation [20-22]. However, this potentially represents a negative feedback loop preventing additional activation of NNMT in skeletal muscle when MeNAM concentrations are high. The chronic effects of NR on skeletal muscle NNMT content and activity, and whole-body and skeletal muscle fatty acid metabolism, warrant further investigation.

## Conclusions

NR supplementation (1000 mg·d^-1^) for seven days did not alter substrate metabolism or mitochondrial biogenic signalling in resting or exercised human skeletal muscle. In contrast, NR supplementation did reduce the exercise-induced expression of NNMT mRNA in skeletal muscle, an enzyme proposed to play a putative role in whole-body fatty acid metabolism. Collectively our data would therefore suggest that NAD^+^ metabolism is tightly regulated in human skeletal muscle, with short-term NAD^+^ precursor supplementation unable to modulate this response at rest or during and in recovery from endurance exercise. Our data therefore adds to a growing list of studies suggesting that NR supplementation does not alter mitochondrial biogenic signalling in healthy human skeletal muscle.

## Acknowledgements

The authors would like to thank the participants for their efforts, their time and their tissue. The authors would also like to acknowledge the support of Lauren Homer and Nathan Hodson in the data collection of this study.

